# Synthesis, SAR and Docking Studies of Substituted Aryl phenylthiazolyl phenylcarboxamide as potential PTP1B Inhibitors

**DOI:** 10.1101/472670

**Authors:** Kanika Varshney, Amit K. Gupta, Arun Rawat, Rohit Srivastava, Akansha Mishra, Mridula Saxena, Arvind K. Srivastava, Sudha Jain, Anil K. Saxena

**Affiliations:** Medicinal & Process Chemistry Division, Central Drug Research Institute, Lucknow, 226031, India; Department of Integrative Biology and Pharmacology, McGovern Medical School, University of Texas Health Science Center at Houston, Houston, Texas 77030, USA; Biochemistry Division, Central Drug Research Institute, Lucknow, 226031, India; Department of Chemistry, Amity University, Lucknow, 226010, India; Department of Chemistry, Lucknow University, Lucknow, 226007, India

**Keywords:** Type 2 diabetes, PTP1B, aryl thiazolyl phenyl benzamide, docking

## Abstract

In our continued effort to discover novel PTP1B inhibitor with improved *in vivo* activity we attempeted to optimize our previously discoved lead compound by replacing the sulfonyl group with benzoyl group to yield compound **II**. Additional structural modifications were performed on compound **II** to yield a series of 24 aryl phenylthiazolyl phenylcarboxamides as potential PTP1B inhibitors. Among these compounds six compounds showed good PTP1B inhibitory activity in the order of compound **38 > 30 > 29 > 37 > 22 > 19**. The plausible PTP1B binding site interaction of compound **38** showed favorable binding similar to known PTP1B binders and suggest its selectivity towards PTP1B. Compound **38** also showed promising antihyperglycemic, antidyslipidemic and insulin resistant reversal activities *in vivo* in STZ model and db/db mice model. Altogether, the compound **38** present an excellent candidate for future PTP1B targeted drug discovery.

## 1. Introduction

Protein-tyrosine phosphatase 1B (PTP1B) is a validated molecular target for the development of novel insulin-sensitizer agents addressing both Type 2 diabetes and obesity. PTP1B catalyses dephosphorylation of the activated insulin receptor (IR) as well as Janus kinase 2 (JAK2) and thus negatively regulate insulin and leptin signalling pathways.^[1–3]^ All of PTPs share a high degree of structural similarity in the active site, pTyr (phosphotyrosine)-binding pocket which poses a chanlenge to design PTP1B selective inhibitors.^[4]^ In 2001, Shen et al. successfully exploited the presence of a secondary PTP1B specific noncatalytic, arylphosphate binding site located close to the active site to design selective bidentate PTP1B inhibitors that can simultaneously occupy the active site and the nearby noncatalytic site.^[5]^ Since then number of PTP1B inhibitors targeting both of these binding sites are Benzooxathiazole derivatives^[6]^, Isothiazolidinone^[7]^, Hydroxyphenylazole derivatives^[8]^, thiazolidine-2,4-dione derivative^[9, 10]^, Thiazolyl derivatives^[11]^ have been reported. These inhibitors although show high PTP1B binding affinity but contain less cellular potency because of poor membrane permeability which is associated with targeting active site by highly acidic phosphotyrosine mimetics.^[1]^ In past we have synthesized novel substituted aryl thiazolyl phenylsulphonamides as nonphosphorous small molecule inhibitors of PTP1B using fragment-based approach, where compound **I** (73.6% PTP1B inhibition at 10µM) was found to be the most active PTP1B inhibitor among the series (Fig. 1).^[12]^ In continuation of our efforts to design more in vivo effective PTP1B inhibitors we herein report further lead optimization of compound **I** using sub-structural approach to identify PTP1B inhibitors with better in vitro and in vivo activities.

**Figure 1.**
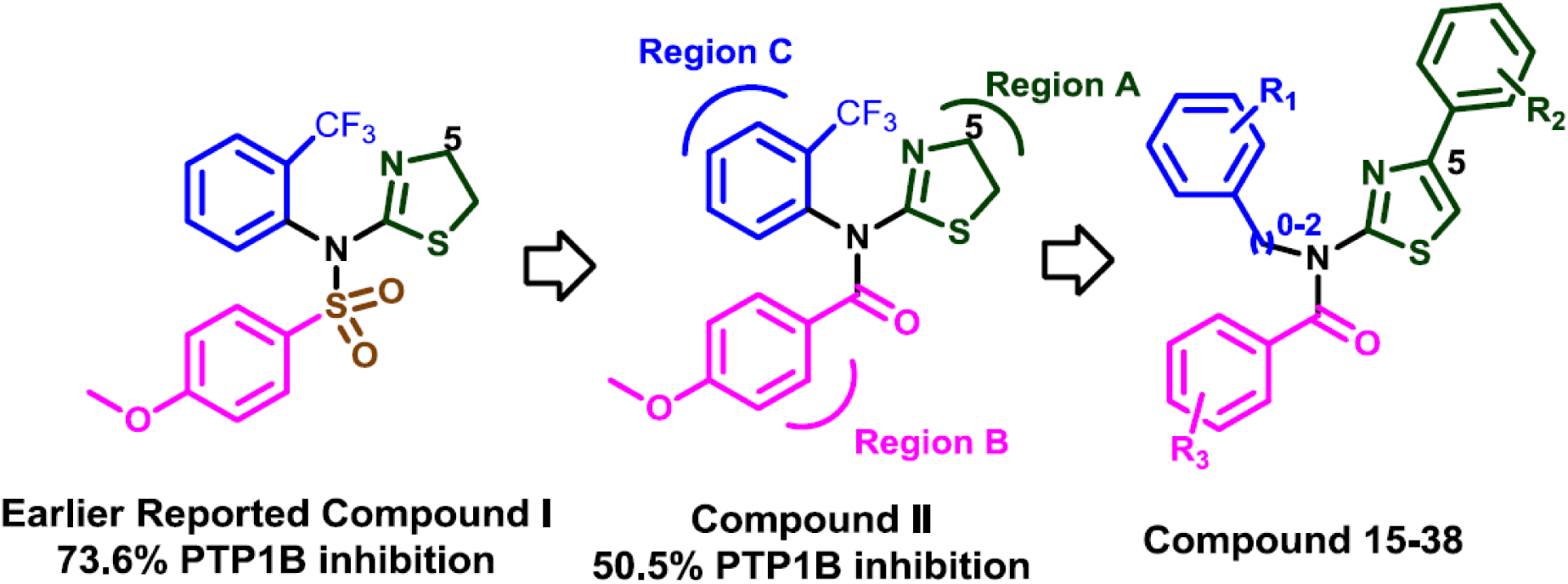
Optimization of compound 1.

## 2. Method and Materials

### 2.1. Chemistry

Melting points were determined on an electrical heated m. p. apparatus /using silicon oil bath. Reactions were monitored by thin layer chromatography on self-made plates of silica gel G (Merck, India) or 0.25mm ready-made plates of silica gel 60F 254, (Merck, Darmstadt, Germany). Column chromatography was performed on silica gel (Merck, 60 to 120mesh). Infrared spectra (IR) were recorded on Perkin-FTIR model PC spectrophotometer with frequency of absorptions reported in wave numbers. Mass were recorded on JEOL mass-spectrometer with fragmentation pattern reported as values, ^1^H NMR was recorded on Bruker spectrometer with a multinuclear inverse probe head with gradient at room temperature (298 K) using CDCl_3_ as solvent and tetramethylsilane (TMS) as internal standard. Chemical shifts were described in parts per million (ppm) relative to TMS (0.00 ppm) using scale and coupling constants were reported in hertz (Hz).

#### General method for synthesis of compound II

The key intermediate N-(2-(trifluoromethyl))-4, 5-dihydrothiazol-2-amine (3d) of compound **II** was synthesized using 2-trifluoromethyl aniline according to the procedure mentioned in the earlier reported paper.^[12]^ Further, the intermediate **3d** on reaction with 4-methoxybenzoylchloride gave the compound **II**.

#### General method for synthesis of compound 7-38

Substituted aniline **(1a-c)**, on reaction with benzoyl isothiocyanate in dry benzene, resulted in the formation of phenylcarbamothioyl benzamides, **(2a-c)** which on alkaline hydrolysis gave the corresponding thioureas **(3a-c)**. Condensation of the phenylthiourea with substituted α-bromo-acetophenones **(4-6)** in THF at room temperature for half-an-hour resulted in the formation of suspension, which was filtered and dried to afford substituted thiazol-2-amines **(7-14)**. These amines on reaction with substituted benzoyl chlorides (R_4_COCl) gave the desired compounds **15-38** (Scheme 1).

**Scheme 1.^a^.**
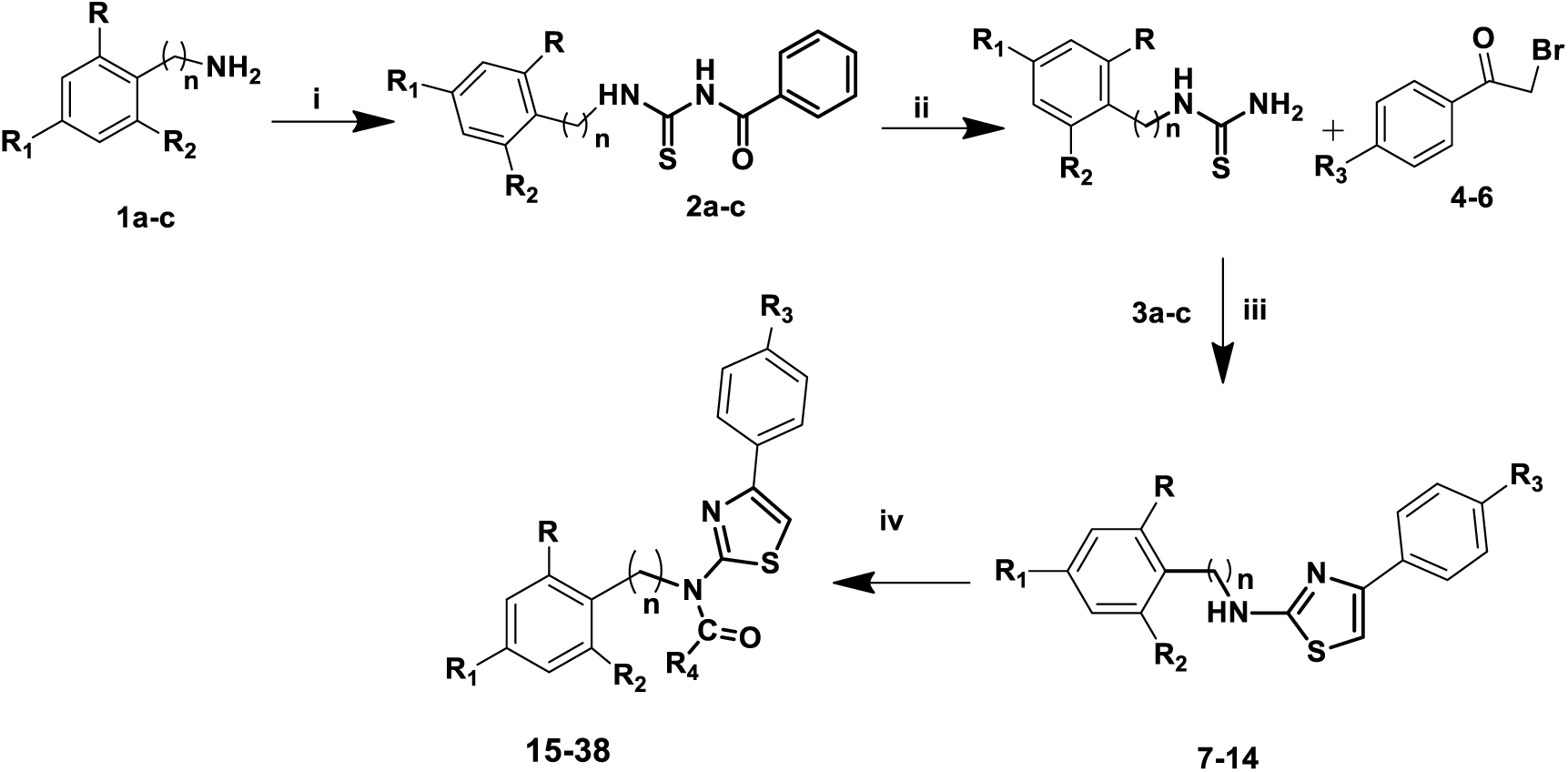
^a^Reagents and conditions (i) C_6_H_5_CONCS, dry benzene (ii) 10 % NaOH (iii)THF,r.t (iv) R_4_COCl, K_2_CO_3_, dry THF, reflux.

### 2.2. *In vitro* PTP1B enzyme assay

All the synthesized compounds were evaluated for *in vitro* antihyperglycemic activity against protein tyrosine phosphatase 1B using colorimetric, non-radioactive PTP1B tyrosine phosphatase drug discovery kit BML-AK822 from Enzo Life Sciences, USA. PTP1B enzyme inhibitory activity of compounds was evaluated using human recombinant PTP1B enzyme provided in the kit at five different concentration i.e. 0.3 µM, 1.0 µM, 3.0 µM, 5.0 µM and 10 µM concentration taking suramin as a control and IC_50_ was calculated for the compounds showing >50% inhibition at 10µM concentration. Other components of the kit include substrate (IR5 insulin receptor residues), biomol red (phosphate determining reagent), assay buffer, suramin (PTP1B inhibitor) and calibration standards. Assay was done according to the Kit manufacturer’s protocol, in brief the reaction was carried out in 96 well flat-bottomed microtiter plate by the addition of assay buffer, solution of test compounds and diluted PTP1B enzyme. Enzyme reaction was initiated by addition of 50µl of warmed 2x substrate then incubated the plate at 300c for 30min. After incubation for 30 min. Reaction was terminated by addition of 25 µl of biomol red reagent and mixed thoroughly by repeated pipetting. Test compounds were dissolved in dimethyl sulfoxide (DMSO) and solution of 100µM concentration was prepared of which 10 µl solution was added in each reaction well to achieve final concentration of 10 µM in reaction mixture. Volumes and dilution of other component were accordingly as instructed in the manual provided in the kit. PTP1B phosphatase acting on the phosphopeptide substrate and release phosphate. The detection of free phosphate released is based on classic Malachite green assay ^[25]^. After adding biomol red to reaction wells after 30 minutes of incubation as described earlier the plate was incubated for another 20 min to develop the colour. Absorbance was recorded at 620 nm on a microplate reader. The percentage inhibition of PTP1b enzyme by test compounds was calculated based on activity in the control tube (without inhibitor) taking as 100 % from three independent set of experiments. The concentration of dimethyl sulfoxide (DMSO) in the test well (1.0 %) had no demonstrable effect on PTP1b enzyme activity.

### 2.3. Molecular docking

To gain an insight about essential structural requirements for ligand interactions at the active site of the PTP1B enzyme, the docking study was performed with the compound **38** at the binding site of PTP1B crystal structure (PDB ID: 2ZMM)^[13]^ using the standard precision (SP) mode of Glide v6.7 of Schrödinger.^[14]^ Ligands were prepared using LigPrep3.4 with Epik3.2 at pH 7.0. The lowest energy conformation among the largest populated cluster sharing common interactions was selected as the best binding pose for docked ligands.^[15]^

### 2.4. *In vivo* asaay

#### 2.4.1. Anti-hyperglycemic activity

Streptozotocin induces anti-hyperglycemic activity screening was performed as reported in ref. 16. Streptozotocin (STZ) is a broad spectrum antibiotic and selected to induce experimental diabetes because of its greater selectivity of β-cells, lower mortality and relatively longer half life (15 min) of STZ in the body. A solution of STZ (60 mg/kg) in 100 mM citrate buffer, pH 4.5 was prepared and calculated amount of the fresh solution dosed to overnight fasted rats intraperitoneally. Two days later baseline blood glucose was drawn from tail vein and glucose levels determined by glucostrips (Roche) to confirm the induction of diabetes. Rats having hyperglycaemia of the range of 270 and 450 mg/dL were considered as diabetic, selected and divided into groups of five animals each. One group used for normal control receives only vehicle (gumacacia) and this group was considered as diabetic control. The blood glucose measured at this time was termed the baseline (0 min) blood glucose. Rats of experimental groups were orally administered suspension of the desired test samples (made in 1.0% gum acacia) at desired dose levels and the biguanide derivative Metformin was used as standard antidiabetic drug and was always given at a dose of 100 mg/kg body weight orally to the experimental group. After 30 min of drug treatment, blood glucose level was again measured with glucometer. The blood glucose assessment were collected from tail vein just prior administration of test sample i.e. 0, 30, 60, 90, 120, 180, 240, 300 and 1440 min post test sample administration. After 300 min the STZ treated animals were allowed to feed over night to overcome drug induced hypoglycaemia. The animals were fed *ad lib* during 5 to 24h of experiments. The average fall in AUC (area under curve) in experimental group compared to control group was termed as % antihyperglycaemic activity. Statistical analysis was done by Dunnett’s test.

#### 2.4.2. db/db mice study

Male C57BL/Ks strain of mouse (db/db mouse) 10-12 weeks of age and around 40 ±3 g of body weight was procured from the animal colony of the Institute. The animals were housed four or five in a polypropylene cage in the animal house. The following norms were always followed for animal room environment: temperature 23 ± 2°C; humidity 50-60%; light 300 lux at floor level with regular 12h light cycle; noise level 50 decibel; ventilation 10-15 air changes per hour. After randomization into groups, the mice were acclimatized for 2-3 days in the new environment before initiation of experiment. Standard pellets were used as a basal diet during the experimental period.

##### 2.4.2.1. Antihyperglycaemic and Antidyslipidemic activity assessment in C57BL/Ks strain of mouse (db/db mouse)

The animals were allocated into groups of 5 animals in each. Prior to start of the sample feeding, a vehicle training period was followed from day −3 to day 0 during which all the animals were given vehicle (1% gum acacia) at a dose volume of 10 mL/kg body weight. At day 0 the animals having blood group level between 350 to 500 mg/dL were selected and divided into three groups containing 5 animals in each. One group was considered as control group while the other group was treatment group. The experimental group was given suspensions of compound **154** and Pioglitazone at 30.0 and 10.0mg/kg body weight dose respectively. The control group was given an equal amount of vehicle. All the animals had free access to fresh water and normal diet. Random blood glucose of each mouse was checked daily at 10.00 pm. On day 10 and day 15 oral glucose tolerance (OGTT) test was performed to study the effect of compound on glucose tolerance. Blood has been withdrawn from the retroorbital plexus of mice eye for the estimation of lipid profile on DIALAB DTN-410-K and insulin level by CALBIOTECH Insulin ELISA Kit. Body weight of each animal was measured on alternate day for studying the effect of test sample on body weight. The skeletal muscle from each mouse were quickly excised at the end of experiment under light anesthesia and frozen at −80°C until further use.

##### 2.4.2.2. Oral glucose and intraperitoneal insulin and pyruvate tolerance test

Oral glucose and pyruvate tolerance test were performed on 12h fasted mice while insulin tolerance test was performed on 4h fasted mice. For OGTT mice were administered glucose 3.0 g/kg by gavages whereas for ITT and PTT, insulin (0.8U/kg) and pyruvate (2g/kg) was injected intraperitoneally, respectively. Blood samples were obtained *via* tail nick at 0, 30, 60, 90 and 120 min during OGTT and PTT while at 0, 15, 30, 60, 90 and 120 time points in ITT. Glucose was measured with the One Touch Ultra glucometer (Accu-Chek Sensor, Roche Diagnostics).

#### 2.4.3. Western blot analysis

Collected tissues and cells were homogenized into PBS containing 1% NP40, 5 mM EDTA, phosphatase inhibitors and protease inhibitors cocktail (lysis buffer). Samples were homogenized and incubated on ice for 15 min. Sample is then stored at – 80°C and thawed at 37°C in waterbath. Sample was then centrifuged at 16000 rpm at 4°C. Then supernatant was taken and quantified by Bradford assay. 40 μg protein (supernatant) of each incubation was resolved on SDS-PAGE, transferred to nitrocellulose membranes and probed with p-Akt and p-IRS1, Glut4 (Cell Signaling, MA, USA) and β-actin (Santa Cruz) was taken as the loading control. Immunoreactive bands were visualized by Enhanced Chemiluminescence according to manufacturer’s instructions (GE Healthcare, UK). Protein expression was evaluated by densitometric analysis performed with Alpha DigiDoc 1201 software (Alpha Innotech Corporation, CA, USA). The same size rectangle box was drawn surrounding each band and the intensity of each was analysed by the program after subtraction of the background intensity.

#### 2.4.4. Statistical analysis

The homeostatic model assessment (HOMA) was used to calculate relative insulin resistance as follows: [Fasting blood glucose (mg/dL) × Fasting serum insulin (µLU/mL)]/405. All statistical calculations were performed using Graph-Pad Prism version 3.02 for Windows (GraphPad Software). Statistical analysis was carried out by Students t test. Data was expressed as mean +SE. The criterion for statistical analysis was significant (*p<0.05), more significant (**p<0.01), highly significant (**p<0.01) and not significant (ns). The results are reported as mean values ± SEM.

## 3. Result and Discussion

### 3.1. Design and synthesis

PTP1B inhibitors containing thiazolyl and carboxyl group are known.^[7, 9–11, 13]^ Therefore, we replaced sulfonyl moiety in the compound **I** by benzoyl group (region B) to afford compound **II** and to see its impact on the PTP1B inhibitory activity. (Fig. 1)

In the next step the dihydrothiazolyl moiety in compound **II** (Region A) was substituted by 5-phenylthiazolyl group to investigate if the phenyl ring can occupy better space/ interactions at the PTP1B enzyme active site than compound II. The systematic substitution by different groups around region B and C was carried out for getting more insight for SAR. This led us to synthesize a series of 24 substituted thiazolyl-N-phenyl-benzamide derivatives using procedure described in scheme 1.

These synthesized compounds were evaluated for PTP1B enzyme inhibitory activity. Compounds more than 50% inhibition at 10 µM concentration were further evaluated at five different concentrations to calculate their IC_50_ values (Table 1). Suramin (supplied in the enzyme activity kit) was used as a reference standard. The antihyperglycemic activity including PTP1B inhibitory activities of the thiazolyl-N-phenylcarboxamide derivatives are also presented in Table 1.

**Table 1:** *In vitro* PTP1B enzyme inhibitory and *in vivo* antihyperglycemic activity in STZ model for compound 15-38.

| Compd. No | R | R<sub>1</sub> | R<sub>2</sub> | R<sub>3</sub> | R<sub>4</sub> | n | % inhibition at 10 μM | IC<sub>50</sub> (μM) | Blood glucose level <sup>b</sup> | |
| --- | --- | --- | --- | --- | --- | --- | --- | --- | --- | --- |
|  |  |  |  |  |  |  |  |  | 0-5hr | 0-24hr |
| II | CF<sub>3</sub> | H | H | - | 4-OMePh | - | 48 |  |  |  |
| 15 | H | F | H | Me | 4-ClPh | 0 | 35.2 (± 3.4) | | | |
| 16 | H | F | H | Me | -CH<sub>2</sub>CH<sub>2</sub>Ph | 0 | 33.0 (± 4.6) | | | |
| 17 | H | F | H | Me | -CH<sub>2</sub>Ph | 0 | 60.3 (± 5.8) | 8.0 (± 2.8) | 14.1 (± 4.7) | 23.2 (± 5.5) |
| 18 | H | F | H | OMe | 4-ClPh | 0 | 56.9 (± 3.8) | 8.8 (± 2.2) | | |
| 19 | H | F | H | OMe | -CH<sub>2</sub>CH<sub>2</sub>Ph | 0 | 74.7 (± 6.7) | 7.0 (± 2.4) | | |
| 20 | H | F | H | OMe | -CH<sub>2</sub>Ph | 0 | 9.8 (± 3.7) | | | |
| 21 | H | F | H | H | 4-ClPh | 0 | 37.2 (± 9.7) | | | |
| 22 | H | F | H | H | -CH<sub>2</sub>CH<sub>2</sub>Ph | 0 | 69.0 (± 5.8) | 6.9 (± 2.1) | 2.44 (± 1.8) | 11.7 (± 4.4) |
| 23 | H | F | H | H | -CH<sub>2</sub>Ph | 0 | 24.7 (± 4.7) | | | |
| 24 | OMe | H | OMe | Me | 4-ClPh | 0 | 26.5 (± 4.2) | | | |
| 25 | OMe | H | OMe | Me | -CH<sub>2</sub>CH<sub>2</sub>Ph | 0 | 56.2 (± 6.2) | 8.6 (± 2.1) | | |
| 26 | OMe | H | OMe | Me | -CH<sub>2</sub>Ph | 0 | 63.6 (± 6.8) | 7.7 (± 1.5) | 11.4 (± 3.6) | 15.8 (± 4.2) |
| 27 | OMe | H | OMe | OMe | 4-ClPh | 0 | 64.1 (± 7.7) | 7.6 (± 1.7) | 13.4 (± 3.8) | 17.8 (± 3.9) |
| 28 | OMe | H | OMe | OMe | -CH<sub>2</sub>CH<sub>2</sub>Ph | 0 | 64.3 (± 7.9) | 7.4 (± 2.8) | 10.6 (± 2.5) | 9.56 (± 2.8) |
| 29 | OMe | H | OMe | OMe | -CH<sub>2</sub>Ph | 0 | 75.0 (± 8.8) | 6.3 (± 2.1) | 17.6 (± 1.9) | 25.1* (± 2.8) |
| 30 | OMe | H | OMe | H | 4-ClPh | 0 | 71.3 (± 9.2) | 7.3 (± 3.5) | 2.48 (± 1.8) | 6.78 (± 2.7) |
| 31 | OMe | H | OMe | H | -CH<sub>2</sub>CH<sub>2</sub>Ph | 0 | 52.7 (± 6.6) | 8.6 (± 4.5) | - | |
| 32 | OMe | H | OMe | H | -CH<sub>2</sub>Ph | 0 | 57.2 (± 8.6) | 8.0 (± 3.8) | 18.3 (± 4.1) | 16.6 (± 4.8) |
| 33 | H | OMe | H | Me | 4-ClPh | 2 | 9.8 (± 3.8) | | | |
| 34 | H | OMe | H | Me | -CH<sub>2</sub>CH<sub>2</sub>Ph | 2 | 24.6 (± 4.2) | | | |
| 35 | H | OMe | H | Me | -CH<sub>2</sub>Ph | 2 | 21.6 (± 4.8) | | | |
| 36 | H | OMe | H | OMe | 4-ClPh | 2 | 14.6 (± 3.7) | | | |
| 37 | H | OMe | H | OMe | -CH<sub>2</sub>CH<sub>2</sub>Ph | 2 | 69.9 (± 5.8) | 6.9 (± 2.7) | 6.10 (± 2.8) | 13.2 (± 4.7) |
| 38 | H | OMe | H | OMe | -CH<sub>2</sub>Ph | 2 | 76.6 (± 7.6) | 5.8 (± 1.8) | 22.8 (± 3.7)* | 23.7 (± 4.8) ** |
| Suramin | - | - | - | - | - | - | 67.3 (± 6.8) | | | |
| Sodium orthovanadate | - | - | - | - | - | - | -- | | 24.3* (± 3.4) | 30.1 (± 4.8) ** |

Values are mean ± SEM, n=4-6, p< 0.05 \* p<0.01\*\*, <sup>b</sup> % lowering at the dose of 100 mg/kg)

### 3.2. Structure activity relationship (SAR) of compounds tested

Initial SAR studies were focused on C5 position of thiazolyl ring (R_3_) to explore if binding with the non-catalytic binding site of PTP1B could be enhanced. Therefore, as a starting point -Me group was inserted at R_3_ position and was kept constant to see the effect of R, R_1_, R_2_ and R_4_ substituent on the PTP1B inhibition.

Compound **17** having R, R_2_ as H, R_1_ as fluoro, R_3_ as Me and R_4_ as CH_2_Ph showed 60.3% PTP1B inhibition (IC_50_ = 8.0 µM). Substitution of R, R_2_ with OMe, R_1_ as H and keeping R_4_ as CH_2_Ph leading to compound **26** resulted in a negligible increase in activity (63.6 % PTP1B inhibition; IC_50_ = 7.7 µM). However, the substitution 2,5-(OMe)_2_Ph (R, R_2_=OMe, R_1_=H) in compound **26** by 4-OMePhCH_2_CH_2_ (R, R_2_=H, R_1_=OMe, n=2) and keeping R_4_ as CH_2_Ph in compound **35** resulted in a great decrease in the activity from 63% to 21.6% PTP1B inhibition.

Further, the replacement of R_4_ group in compound **17** by 4-ClPh (compound **15**) and by CH_2_CH_2_Ph (compound **16**) led to 35% and 33% reduction in PTP1B activity respectively as compared to compound **17 (**60.3% PTP1B inhibition). Similar trends were observed in analogous compounds **24**, **25** with 2,5-(OMe)_2_ (R, R_2_=OMe, R_1_=H) and compounds **33** and **34** with 4-Me (R, R_2_=H, R_1_=OMe, n=2) substitutions. To further explore the SAR, Me group at R_3_ position in the above compounds was replaced by OMe group. In general, the compounds with OMe at R_3_ position were more active in comparison to the compounds with Me group except the compound **20** and **36** which showed less than 15% PTP1B inhibition. The compound **38** (R, R_2_=H, R_1,_ R_3_=OMe, n=2, R_4_=CH_2_Ph) was found to be the most active compound of the series with 76.6% PTP1B inhibition (IC_50_ = 5.8 µM). The replacement of R, R_1_ and R_2_ in compound **38** by R, R_2_ =OMe, R_1_=H in the compound **29** resulted into negligible change in activity (75% PTP1B inhibition at 10 µM; IC_50_=6.3 µM) compared to compound **38**. However, insertion of the F group at R_1_ in place of H and R, R_2_=H in place of OMe led to the substantial decrease in the activity and made compound **20** as least active compound of the series (9.8% PTP1B inhibition). Further, the replacement of CH_2_Ph group at R_4_ position by CH_2_CH_2_Ph resulted in slight reduction in the activity as the compound **37** (R, R_2_=H, R_1_=OMe, n=2, R_3_= OMe, R_4_=CH_2_CH_2_Ph, 69.9% PTP1B inhibition; IC_50_=6.9 µM); and **28** (R, R_2_=OMe, R_1_=H, R_3_=OMe, 64.3% PTP1B inhibition;IC_50_=7.34 µM);. In contrast to the compounds **37** and **28**, the compound **19** (R, R_2_=H, R_1_=F, R_3_=OMe and R_4_=CH_2_CH_2_Ph) showed increased activity with 74.7% PTP1B inhibition (IC_50_=7.0 µM). Incorporation of 4-ClPh group in the place of CH_2_CH_2_Ph at R_4_ position in compound 18 (R, R_2_=H, R_1_=F, R_3_=OMe) showed decrease in activity with 56.9% PTP1B inhibition (IC_50_=8.6µM);. A similar trend was observed in case of compound **27** (R, R_2_=OMe, R_1_=H, R_3_=OMe, 64.1% PTP1B inhibition; IC_50_=7.55 µM); but in compound **36** (R, R_2_=H, R_1_=OMe, n=2, R_3_=OMe) which showed highly reduced activity with 14.6% PTP1B inhibition in comparison to its analogous compound. Removal of the 4-substituent at R_3_ position and keeping it as H (compound nos. **21**, **31** and **32**) resulted in slight decrease in the PTP1B inhibitory activity in comparison to their corresponding 4-OMe and 4-Me analogues except compounds **22** and **30** which showed slight increase in activity in comparison to corresponding analogues with 69% PTP1B inhibition IC_50_=6.9 µM); and 71.3% PTP1B inhibition (IC_50_ = 7.33 µM) respectively.

### 3.3. Kinetics Measurements and Mechanism of Inhibition

Kinetic study was done to determine the type of inhibition of the compound **38**. Enzyme activity assay at different concentration of of compound **38** (0, 5, 10, and 20µM) with varying concentration (40 to 140) µM of substrate IR5 (sequence from the insulin receptor ß subunit domain provided in the kit) was performed and Lineweaver-Burk double reciprocal plot was drawn. (Fig 2)

**Figure 2.**
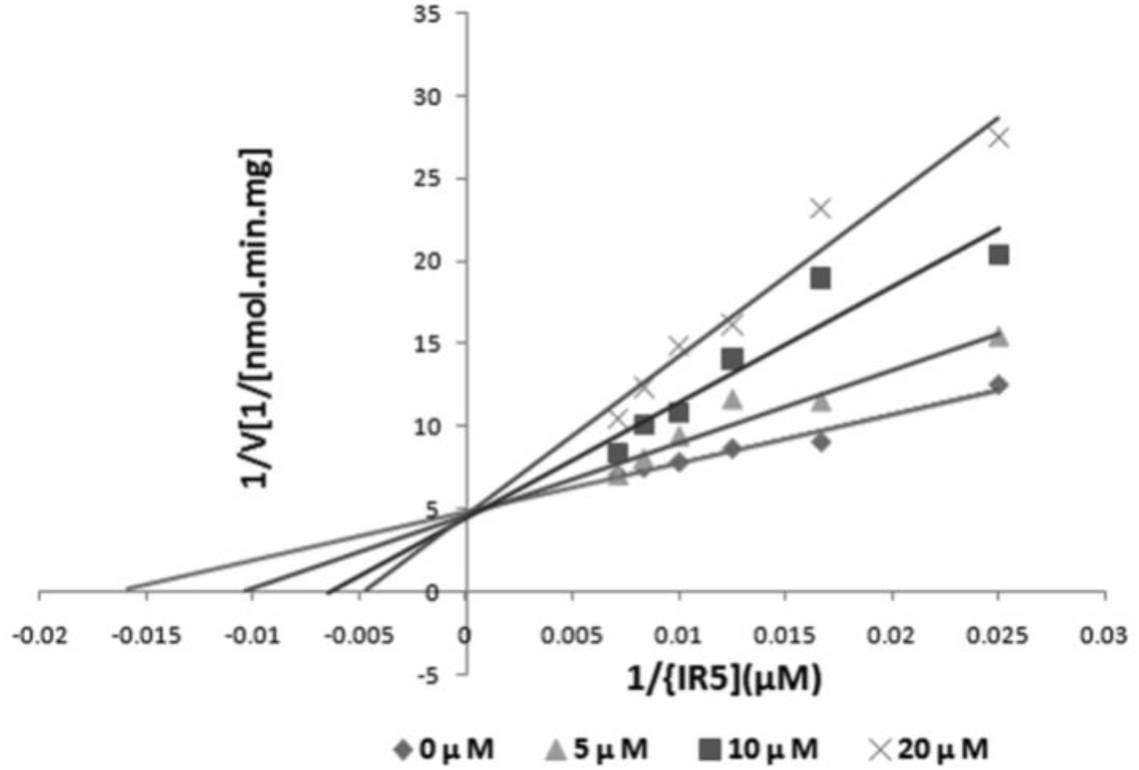
Competitive inhibitory profile of compound **38**.

Fig. 2 Plot shows intercept of all lines obtained with 0µM, 5µM, 10µM and 20µM compound concentration converging at Y-axis (1/V_max_) where slope and X-axis intercept (−1/Km) vary with inhibitor concentration. This suggests compound **38** follows a competitive inhibition kinetics. The K_m_ value for the substrate IR5 was calculated from this plot was found to be 60.1µM. The value of Ki 8.59 µM was calculated from intercept on X-axis of this replot which was determined by plotting the slope values vs compound **38** concentration.

### 3.4. Molecular Modeling Studies

In past our lab has successfully exploited structure-based computational approaches for understanding the structure-activity relationships.^[13, 17–20]^ To gain structural insight for further optimization, we constructed the 3D binding model of compound **38** to PTP1B using molecular docking. The binding analysis of the compound **38** revealed that 4-methoxyphenyl group linked with thiazolyl moeity was buried deep inside of a hydrophobic cavity formed by the residues Phe182, Ile219, and Ala217 which was further stabilized by its π-π interaction with Phe182, which in turn involved in the WPD loop closure. The thiazolyl group itself made π-π interaction with Tyr46 (Fig. 3) which is commonly observed in many known PTP1B/inhibitor complexes.^[14, 21–24]^ The amino acid residues Arg47 and Asp48 which are close to the active site have been demonstrated important for selectivity of PTP1B because of the point mutation at this residue present in other phosphatases such as LAR (Asn48) and SHP-2 (Asn48).^[25, 26]^ The phenyl acetyl group in the compound **38** showed potential to form hydrogen bond interaction with Asp48 and was in close contact to Arg47 which suggest its selectivity towards PTP1B. However, more experiments need to be done to confirm this hypothesis. Water molecules also plays an important role in the stabilization of compound **38** through water mediated hydrogen bond interactions with amino acid residue Asp181 which is consistent with co-crystallized ligand used in this study.^[14]^ Because of these multiple interactions, compound **38** positioned itself effectively at the PTP1B binding site to serve as a potent inhibitor.

**Figure 3.**
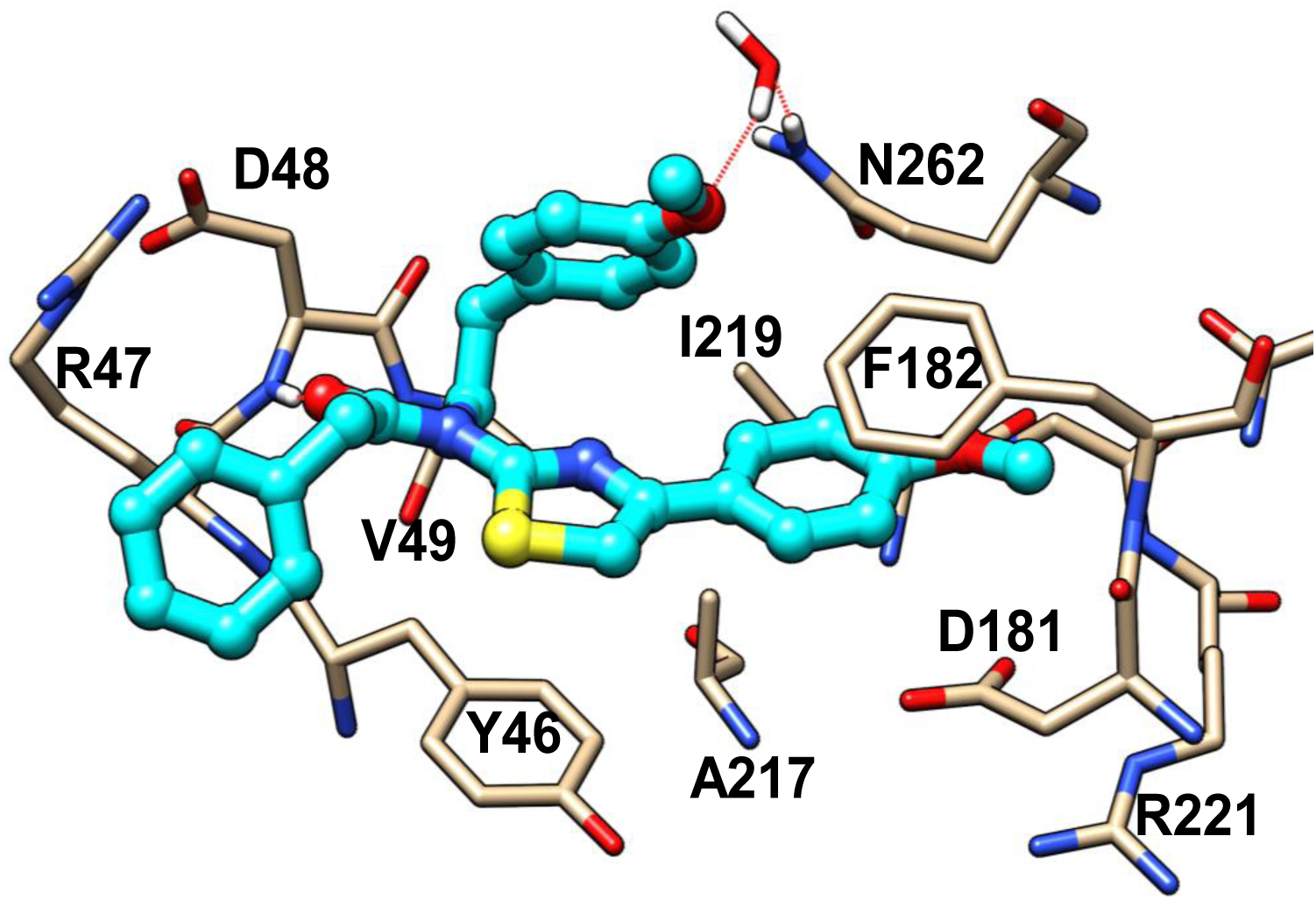
Lowest energy docking pose of compound **38** (cyan) at the active site of PTP1B (PDB ID: 2ZMM).

### 3.5. *In vivo* biological activity

#### 3.5.1. Streptozotocin rat model study

Based on the high PTP1B inhibitory activity *in vitro*, a limited number of compounds with more than 57% PTP1B inhibition were evaluated *in vivo* in Streptozotocin induced rat model (STZ). The table 1 discribes the effect of the ten selected compounds **17**, **22**, **26-30**, **32**, **37**, **38** along with the standard compound Sodium orthovanadate on decline in blood glucose level of streptozotocin treated diabetic rats. Sodium orthovanadate was taken as positive control. It is evident from the results that out of the ten compounds tested, only one compound **38** showed a significant decline in blood glucose levels on streptozotocin induced diabetic rats. This decline in blood glucose level was around 22.8 % (p<0.05) and 23.7 % (p<0.01) during 0-5h and 0-24h, respectively (Fig. 4A and B). The standard drug sodium orthovanadate demonstrated maximum decline in blood glucose levels to the tune of 24.3 % (p<0.05) during 0-5h and 30.1 % (p<0.01) during 0-24h, post treatment respectively on STZ-induced diabetic rats at 100 mg/kg oral dose. The compounds **17**, **22**, **26-30**, **32** and **37** though showed decline in blood glucose on STZ-induced diabetic rats at 100 mg/kg oral dose level as shown in Table 1, however the effect was not found statistically significant in any of the cases except the compound **38** where the decline in blood glucose was observed around 23.7 % during 0-24 h (Fig. 4).

**Figure 4.**
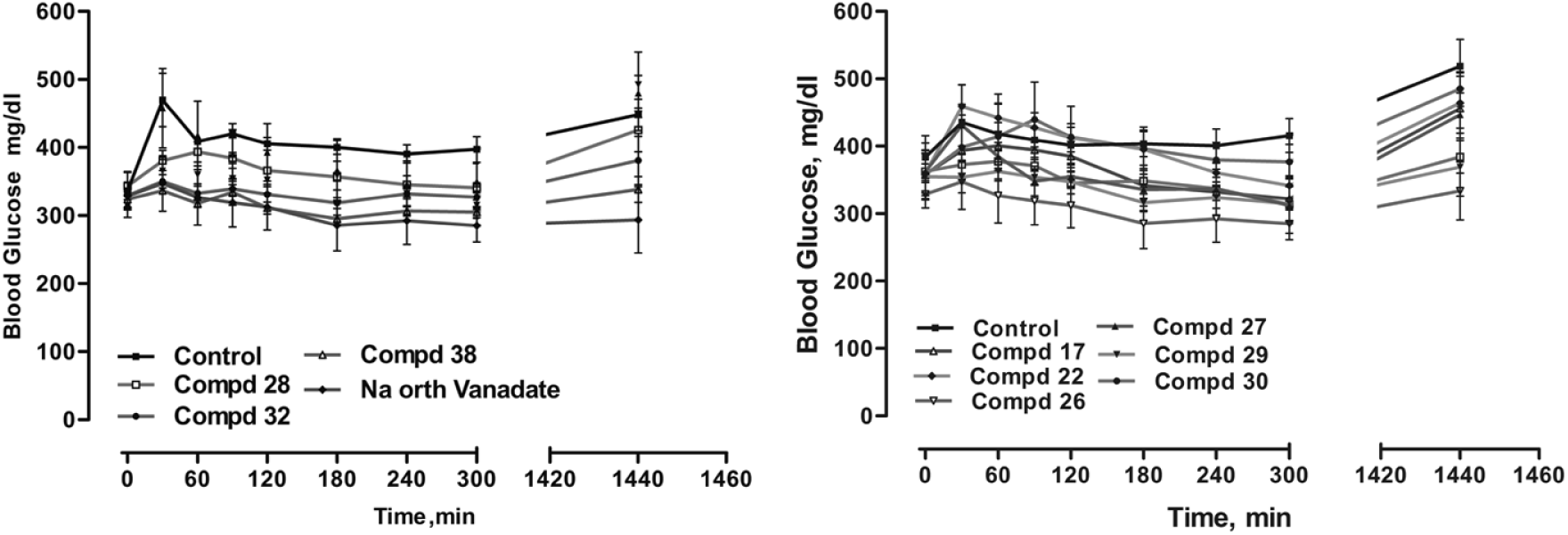
**A** and **B.** Effect of compound **17, 22**, **26-30**, **32, 38** and Standard compound Sodium orthovanadate on blood glucose levels of the streptozotocin treated diabetic rats at various time intervals.

#### 3.5.2. db/db mouse model study

##### 3.5.2.1. Effect on hyperglycaemia

Compound **38** was further evaluated for antihyperglycaemic and antidyslipidemic activities in C57BL/KsJ-db/db mice (Table 2). Antihyperglycaemic activity was carried out *in vivo* using db/db mice by observing overall glucose lowering effect and also improvement on oral glucose tolerance at 30mg/kg for the period of 15 days. Pioglitazone was taken as positive control. It is evident from the Fig. 5A that compound **38** exerted its effect on blood glucose from day 5 while the significant effect was observed from day 7 and persisted till the end of the experiment, whereas, Pioglitazone (at the dose of 10mg/kg) significantly declined the random blood glucose from day 6 which persisted up to the end of the experiment as compared to vehicle treated control group.

**Figure 5:**
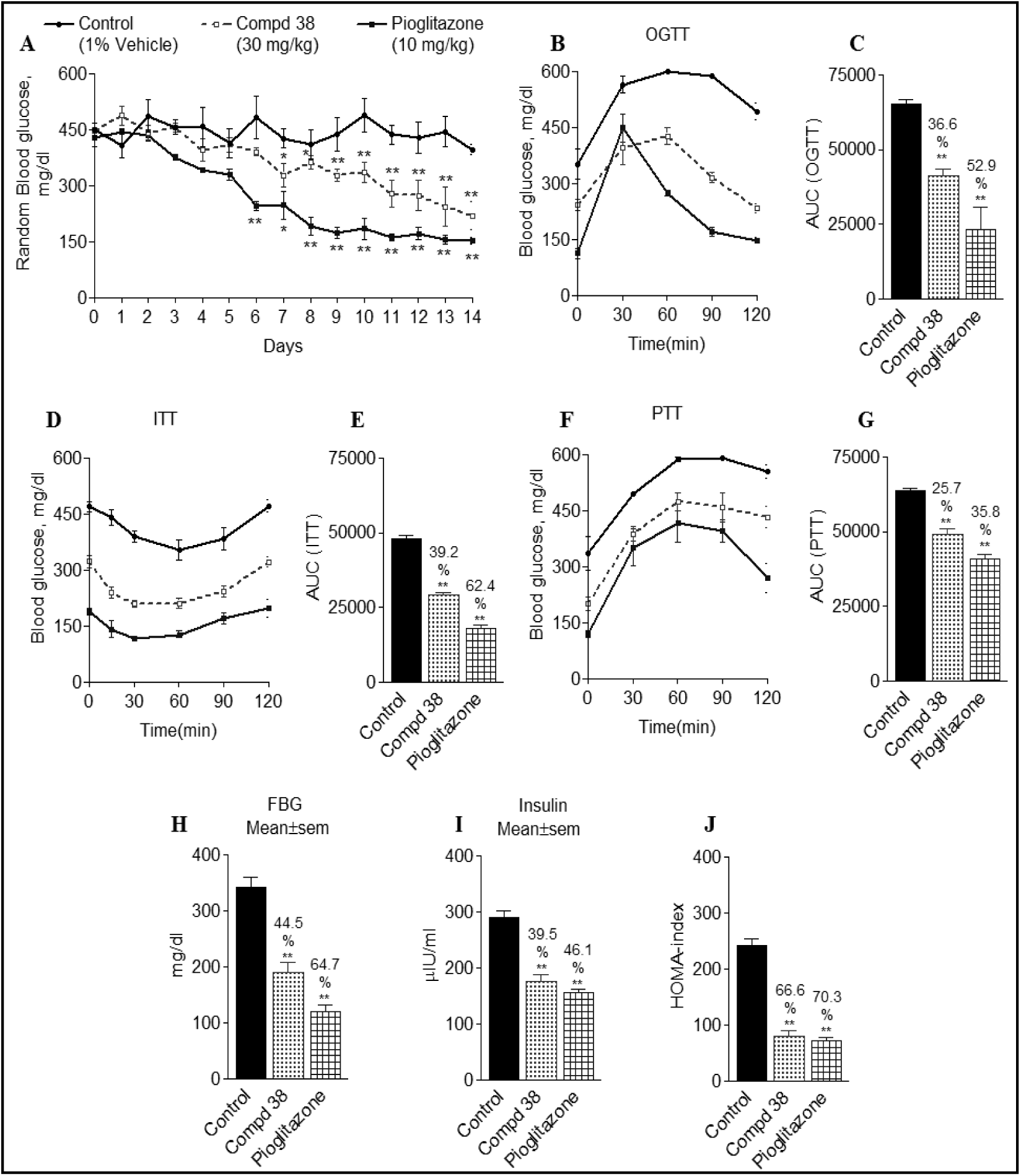
Antihyperglycaemic profile and insulin resistance reversal profile of db/db mice; **A.** Random blood glucose, **B.** Oral glucose tolerance, **C.** Glucose AUC (OGTT), **D.** ITT, **E.** AUC (ITT), **F.** PTT, **G.** AUC (PTT), **H.** FBG, **I.** serum Insulin, **J.** HOMA-index.

**Table 2:** Antihyperglycaemic and antidyslipidemic activity in db/db mice (% efficacy after 15 days)

| Comp<br>. no. | Antihyperglycemic activity |  |  | Antidyslipidemic activity |  |  | Insulin resistant reversal |  |  |
| --- | --- | --- | --- | --- | --- | --- | --- | --- | --- |
|  | OGTT<br>15 days | ITT | PTT | TG | CHOL | HDL | Fasting<br>blood<br>glucose | Serum<br>insulin<br>level | HOMA index |
| **38** | **36.6 ± 3.8** | **39.2 ± 1.9** | **25.7 ± 1.9** | **18.5 ± 0.6** | **11.6 ± 0.3** | **27.3 ± 0.3** | **44.5 ± 5.4** | **39.4 ± 3.4** | **66.6 ± 11.6** |
| Pioglitazone | 52.9 ± 17.6 | 62.4 ± 7.3 | 35.8 ± 2.1 | 12.6 ± 0.6 | 9.2 ± 0.8 | 11.1 ± 0.8 | 64.7 ± 6.6 | 46.1 ± 1.8 | 70.3 ± 10.6 |

##### 3.5.2.2. Effect on oral glucose tolerance

To observe the effect of drug on glucose tolerance, an OGTT (oral glucose tolerance test) conducted on day 14 during dosing in overnight fasted mice. After 14 days of consecutive dosing an OGTT was carried out and the results showed that inhibition of PTP1B by the compound **38** effectively resisted the rise in postprandial hyperglycaemia during 30 and 60 mins post glucose load and significantly enhanced the glucose clearance from blood at 60, 90 and 120 time points (Fig. 5B) and an overall improvement of 36.6% (p<0.01) on glucose disposal was calculated by AUC analysis (0-120min) with no observed hyperglycaemia, whereas, reference drug Pioglitazone showed an improvement of 52.9% (p<0.01) on glucose AUC as shown in Fig. 5C. The fasting baseline blood glucose value at 0 time point was also found lower in the compound **38** treated groups compared to the vehicle treated control group at the corresponding time point because of the antihyperglycaemic effect. It is evident from the graph that the compound **38** and Pioglitazone effectively declined the rise in postprandial hyperglycaemia induced by glucose load of 3g/kg.

To assess the whole-body insulin sensitivity, insulin tolerance test (ITT) was performed in vehicle, compound **38** and pioglitazone treated mice. It was found that compound **38** significantly improved the insulin sensitivity at 15 and 30 min after a bolus of 0.75 units of human insulin as shown in Fig. 5D and area under the curve (AUC) analysis also showed a 39.2% improvement in the insulin tolerance (Fig. 5E) as compared to vehicle treated control group, whereas reference drug Pioglitazone improved the insulin sensitivity by 62.4% (p<0.01). Further to assess the effect of the compound **38** on hepatic insulin sensitivity, the pyruvate tolerance test was performed by intraperitoneally administrating pyruvate, a major gluconeogenic substrate. The results of pyruvate tolerance test showed that the compound **38** significantly increased the insulin sensitivity and effectively resisted the rise in blood glucose (Fig. 5F) caused by a bolus of pyruvate injection as compared to vehicle treated control group. The AUC analysis indicated an improvement of 25.7 and 35.8% (p<0.01) (Fig. 5G) on pyruvate tolerance by compound **38** and reference drug Pioglitazone, respectively as compared to vehicle treated control group.

##### 3.5.2.3. Effect on fasting blood glucose, serum insulin level, HOMA-index

Insulin resistance is one of the characteristic features of db/db mice. Decreased insulin sensitivity leads to hyperglycaemia, hyperinsulinemea. Treatment with the compound **38**, nearly normalizes the fasting blood glucose by 44.5% (p<0.01) and restores the altered insulin level by 39.4% (p<0.01) in the treated diabetic mice as compared to vehicle treated control group as shown in Fig. 5H and 5I. Homeostatic model assessment (HOMA) is a method used to quantify insulin resistance that is quantified on the basis of fasting blood glucose and fasting serum insulin level. The treatment with the compound **38** significantly improves the insulin resistance state by improving the HOMA-index by 66.6% (p<0.01) (Fig. 5J).

##### 3.5.2.4. Antidyslipidemic activity

Type 2 diabetes is known to be associated with several adverse cardiovascular risk factors, including obesity, hypertension and serum lipid abnormalities, characterized mainly by elevated serum total triglycerides and low levels of high-density lipoprotein (HDL) cholesterol. Therefore, compound **38** was further tested for its effect on serum lipid profile. The treatment of the compound **38** restored the altered serum lipid profile as evident from the Fig.6 (A, B and C). The compound **38** significantly lowerd the triglyceride and serum cholesterol levels by 18.5% and 11.5 % (p<0.05) respectively and significantly increased theserum HDL-cholesterol by 27.3.8% (p<0.05) in comparison to the standard drug Pioglitazone which significantly lowered the serum triglyceride and cholesterol levels by 12.6% (p<0.05) and 9.7% respectively.

**Figure 6:**
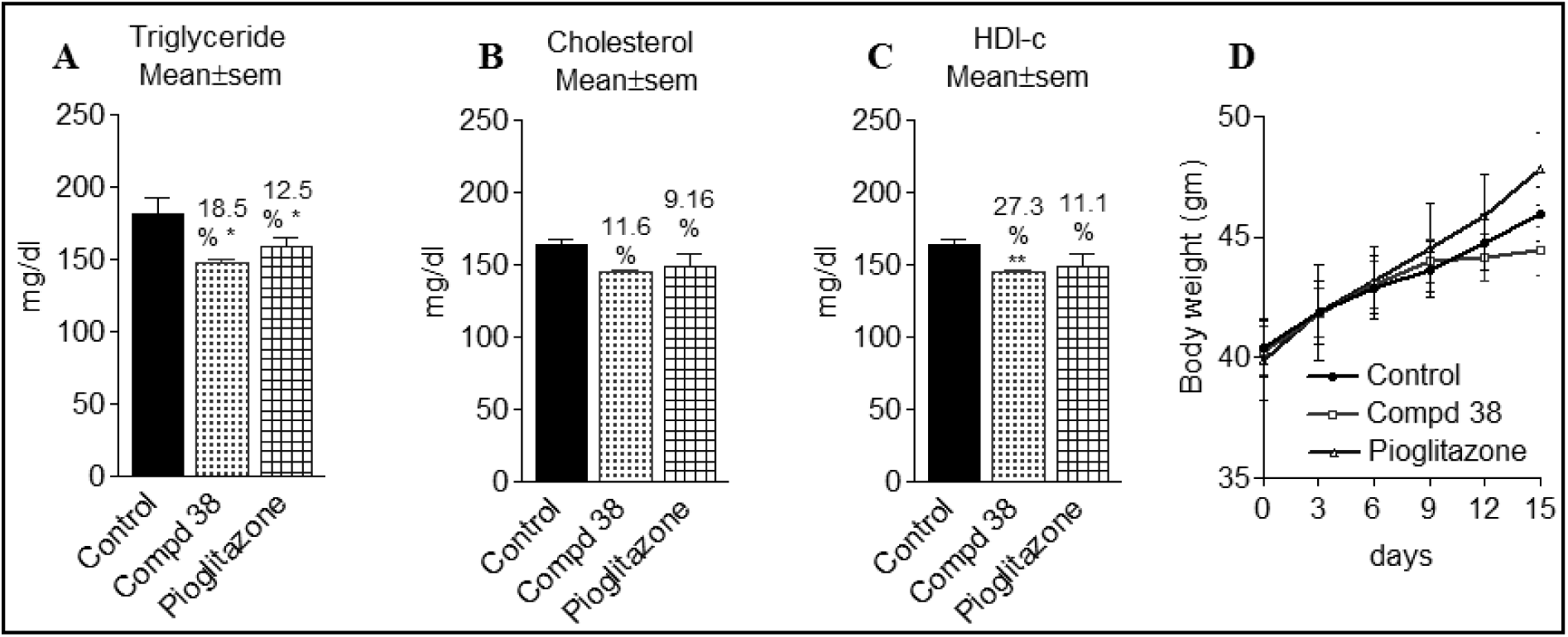
Effect of compound **38** on serum lipid profile of db/db mice, **A)** Triglyceride; **B)** Cholesterol; **C)** HDL-c; **D)** Body weight.

Excessive consumption of the food intake and body weight is responsible for the development of obesity and it can be directly linked to type 2 diabetes. Further observation on body weight during the antidiabetic activity in db/db mice study resulted in decrease of body weight in mice during the treatment of compound **38** (Fig. 6D).

##### 3.3.2.5. Western blot analysis of p-IRSI in skeletal muscle of db/db mice

Further, the inhibition of PTP1B by compound **38** was evident by western blot analysis as shown in Fig. 7A. The compound **38** effectively inhibited the PTP1B which is evident with increase of approximately 2.34 fold increase in the level of p-IRS-1 and 1.95 fold increase in the level of p-Akt, a downstream receptor of insulin signaling, while the reference drug rosiglitazone exhibited an increase of 3.21 and 2.52 folds in p-IRS1 and p-Akt respectively, as compared to vehicle treated control group. (Fig. 7 B and C). The activation of IRS1 and Akt by insulin resulted in the clearance of circulating blood glucose which is evident by the significant increase in protein level of Glut4 in skeletal muscle. In the present study, it was observed that the inhibition of PTP1B with compound **38** resulted in improved insulin signaling and glucose homeostasis with an increase of approximately 2.1-fold in Glut4 (Fig. 7D).

**Figure 7:**
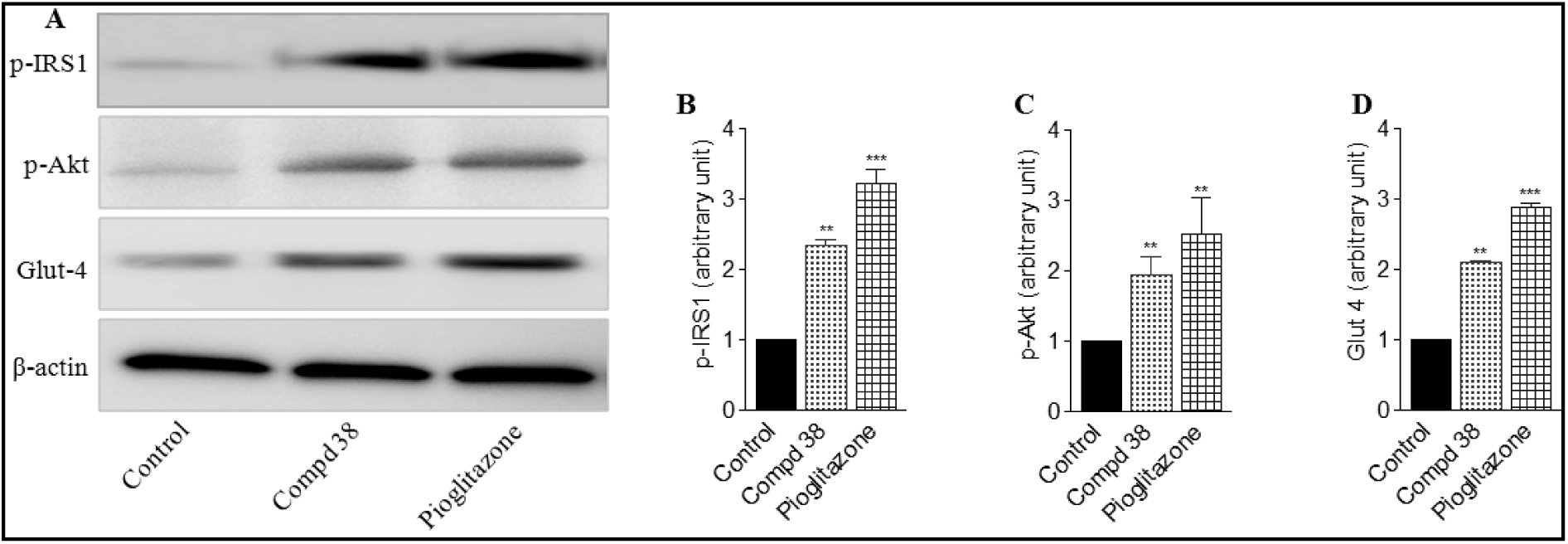
Inhibition of PTP1B by compound **38** improved insulin signaling in skeletal muscle of db/db mice. **A)** Western blot analysis of p-IRSI, p-Akt and Glut4 in skeletal muscle of db/db mice, 40 µg of protein was resolved on SDS-PAGE; **B)** Evaluation for levels of p-IRS1; **C)** Evaluation for levels of p-Akt; **D)** Evaluation for levels of Glut4. The experiments were repeated three times and values are means ± SEM of three independent experiments. The blot shown were representatives of the indicated groups and the densitometric analyses of the same are given below. *p< 0.05, **p < 0.01

## 4. Conclusion

A series of aryl phenylthiazolyl phenylcarboxamide derivatives were synthesized and evaluated against PTP1B enzyme. Among the twenty-five synthesized compounds, six compounds showed good PTP1B inhibitory activity, namely **19** (IC_50_=7.0 μM), **22** (IC_50_=6.9 μM), **29** (IC_50_=6.3 μM), **30** (IC_50_=7.33 μM), **37** (IC_50_=6.9 μM) and **38** (IC_50_=5.8 μM). Docking studies showed that the interacting residues with compound **38** at the active site are consistent with known PTP1B inhibitors. Also interaction of compound **38** with Arg47 and Asp48 may attribute to its selectivity over other homologous phosphatases. Compound **38** also showed promising antihyperglycaemic, antidyslipidemic and insulin resistant reversal activities *in vivo*, in STZ model and db/db mice model. Thus, these studies may be helpful in developing novel PTP1B inhibitors with improved pharmacological properties.

## Acknowledgments

We thank Sophisticated Analytical Instruments Facility, CDRI for providing analytical data. AKG is supported by a fellowship from the Gulf Coast Consortia, on the Computational Cancer Biology Training Program (CPRIT Grant No. RP170593). The CDRI communication number allotted to this manuscript is 01/2019/AKS.

The authors declare no conflict of interests.

## Supporting Information

Method of preparation of intermediate compound 2a, 2b, 3a, 3b and 3c and Characterization data for compound 7-38

